# Significance of trends in gait dynamics

**DOI:** 10.1101/677948

**Authors:** Klaudia Kozlowska, Miroslaw Latka, Bruce J. West

## Abstract

Trends in time series generated by physiological control systems are ubiquitous. Determining whether trends arise from intrinsic system dynamics or originate outside of the system is a fundamental problem of fractal series analysis. In the latter case, it is necessary to filter out the trends before attempting to quantify correlations in the noise (residuals). For over two decades, detrended fluctuation analysis (DFA) has been used to calculate scaling exponents of stride time (ST), stride length (SL), and stride speed (SS) of human gait. Herein, rather than relying on the very specific form of detrending characteristic of DFA, we adopt Multivariate Adaptive Regression Splines (MARS) to explicitly determine trends in spatio-temporal gait parameters during treadmill walking. Then, we use the madogram estimator to calculate the scaling exponent of the corresponding MARS residuals. The durations of ST and SL trends are determined to be independent of treadmill speed and have distributions with exponential tails. At all speeds considered, the trends of ST and SL are strongly correlated and are statistically independent of their corresponding residuals. The group-averaged values of scaling exponents of ST and ST MARS residuals are slightly smaller than 0.5, indicating weak anti-persistence. Thus, contrary to the interpretation prevalent in the literature, the statistical properties of ST and SL time series originate from the superposition of large scale trends and small scale fluctuations. We show that trends serve as the control manifolds about which ST and SL fluctuate. Moreover, the trend speed, defined as the ratio of instantaneous values of SL and ST trends, is tightly controlled about the treadmill speed. The strong coupling between the ST and SL trends ensures that the concomitant changes of their values correspond to movement along the constant speed goal equivalent manifold as postulated by Dingwell et al. doi:10.1371/journal.pcbi.1000856.

**Author summary:** During treadmill walking, the subject’s stride time (ST) and stride length (SL) must yield a stride speed which can fluctuate over a narrow range centered on the treadmill belt’s speed. The fact that both ST and SL are persistent is an intriguing property of human gait. For persistent fluctuations any deviation from the mean value is likely to be followed by a deviation in the same direction. To trace the origin of such persistence, we used a novel approach to determine trends in spatio-temporal gait parameters. We find that the trends of ST and SL of a subject are strongly correlated and are statistically independent of their corresponding residuals. Moreover, the trend speed, defined as the ratio of instantaneous values of SL and ST trends, is tightly controlled about the treadmill speed. The persistence of gait parameters stems from superposition of large scale trends and small scale fluctuations.

## Introduction

Over two decades ago Hausdorff et al. [1, 2] discovered long-range, persistent correlations in stride duration (time) of human gait. Their choice of fractional Brownian motion (FBM) [3] for modelling such correlations has significantly influenced the way in which fluctuations of spatio-temporal gait parameters (stride time, stride length, and stride speed) were subsequently quantified and interpreted. Determining the source of ubiquitous trends observed in physical, social, or biological systems is a recurring problem in fractal time series analysis. In other words, one has to establish whether trends arise from the intrinsic dynamics of a system or have an external origin. In the latter case, a possible approach is first to recognize trends and then filter them out before attempting to quantify correlations in the noise (residuals). In their original study Hausdorff et al. applied detrended fluctuation analysis (DFA) [4] to estimate the scaling (Hurst) exponent which characterizes properties of fractal time series. Their choice of DFA implied that a stride duration time series was made up of persistent (scaling exponent greater than 0.5) fractal fluctuations superposed on trends which are irrelevant from the point of view of fractal analysis.

The papers of Hausdorff et al. spurred significant interest in the emerging field of fractal physiology [5, 6]. In particular, they focused research on the intrinsic variability of physiological time series and their persistence [7]. Some argued that persistent fluctuations are a manifestation of the adaptability of underlying control systems and the absence of long-range correlations, or anti-persistence, indicates disease or pathology [8–10]. Deligni’eres and Torre demonstrated that when healthy humans walk over ground in sync with a metronome, their stride times become anti-persistent [11]. During treadmill walking, a subject’s stride time (ST) and stride length (SL) must yield a stride speed (SS) that fluctuates over only a narrow range centered on the treadmill belt’s speed. It turns out that subjects’ SS is anti-persistent, while the other two gait parameters, ST and SL, are persistent. On the other hand, all three parameters are anti-persistent during treadmill walking with either auditory [12] or visual [13] cueing (alignment of step lengths with markings on the belt). In light of these latter experimental findings, anti-persistence, rather than being a manifestation of pathology, seems to be indicative of tight control.

While persistence of ST and SL during treadmill walking is intriguing, an even more puzzling property of gait dynamics is the weak coupling between stride time and stride length measured by cross-correlation. The coefficient of correlation between these dynamical variables for uncued treadmill walking is 0.28 and increases to approximately 0.55 under the influence of a persistent fractal metronome [14]. Herein, we employ an explicit algorithm to determine trends in spatio-temporal gait parameters and examine their statistical properties as well as those of the corresponding residuals. In the process, light is shed on the the origin of ST and SL persistence.

## Results

### ST and SL trends

The examples of stride time, length and speed time series are given in Fig 1. The solid, thick lines in this figure represent trends approximated using the piecewise linear variant of the MARS model.

**Fig 1.**
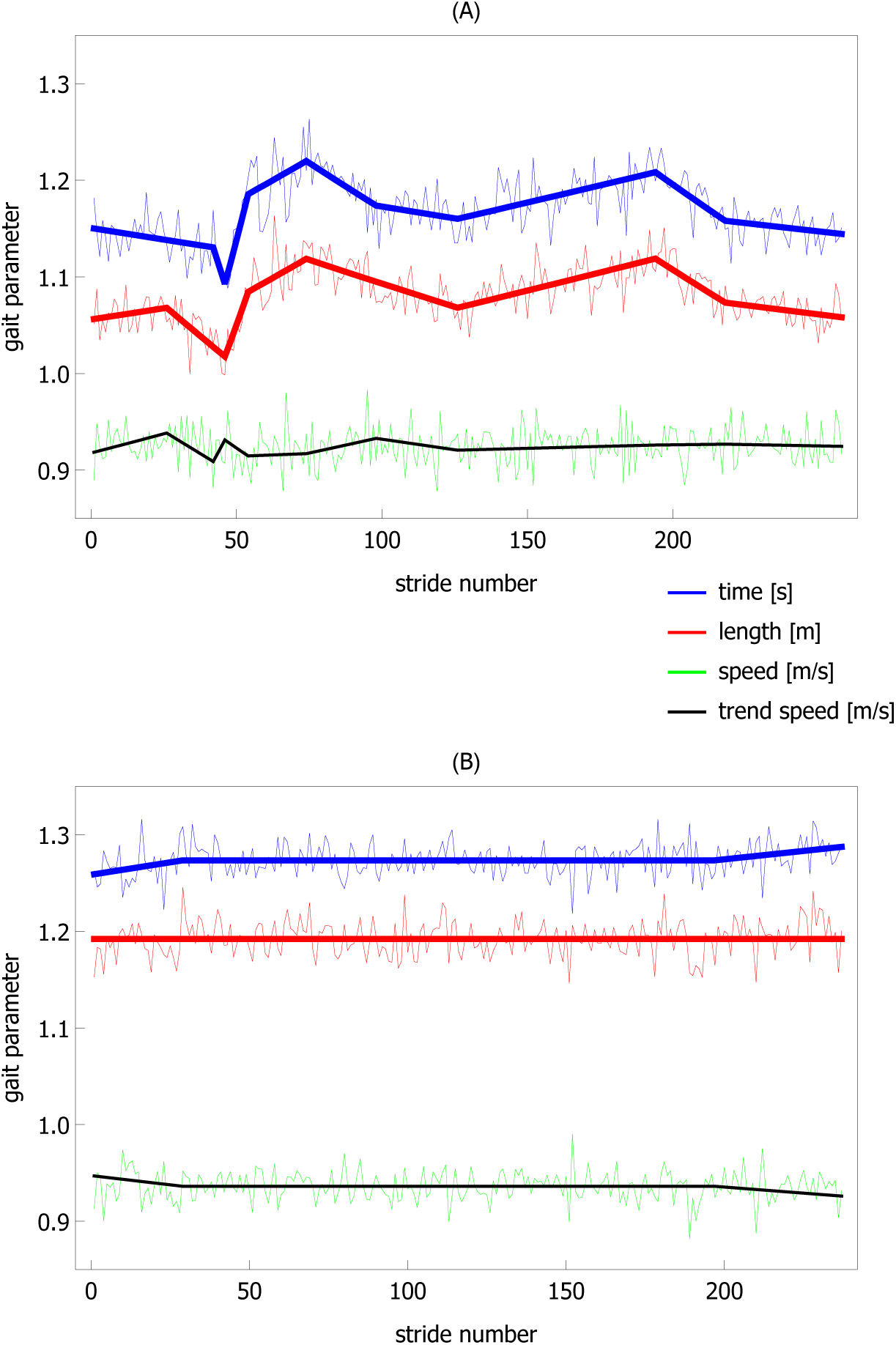
Stride time, length, and speed of two healthy young subjects during treadmill walking at a preferred walking speed: (A) subject 6, trial 2, (B) subject 1, trial 2. Thick solid lines show trends calculated using the piecewise linear variant of the MARS model. The thin solid lines correspond to trend speed (the ratio of values of stride length and stride time trends). Experimental data come from the study of Dingwell et al. [15].

The Kruskal-Wallis test showed that the durations of normalized trends (durations divided by the subject’s mean value of stride time or stride length) were independent of treadmill speed (*p*_*ST*_ = 0.30, *p*_*SL*_ = 0.81). Therefore, we aggregated data from all the trials. In Fig. 2 we present the probability density functions (PDFs) of 1607 ST and 1396 SL trend durations. The Anderson-Darling test showed that the distributions of both gait parameters were not different (*p* = 0.59). The average values and standard deviations of trend durations were close to each other (23.2 and 26.3 for ST, and 25.2 and 35.5 for SL).

**Fig 2.**
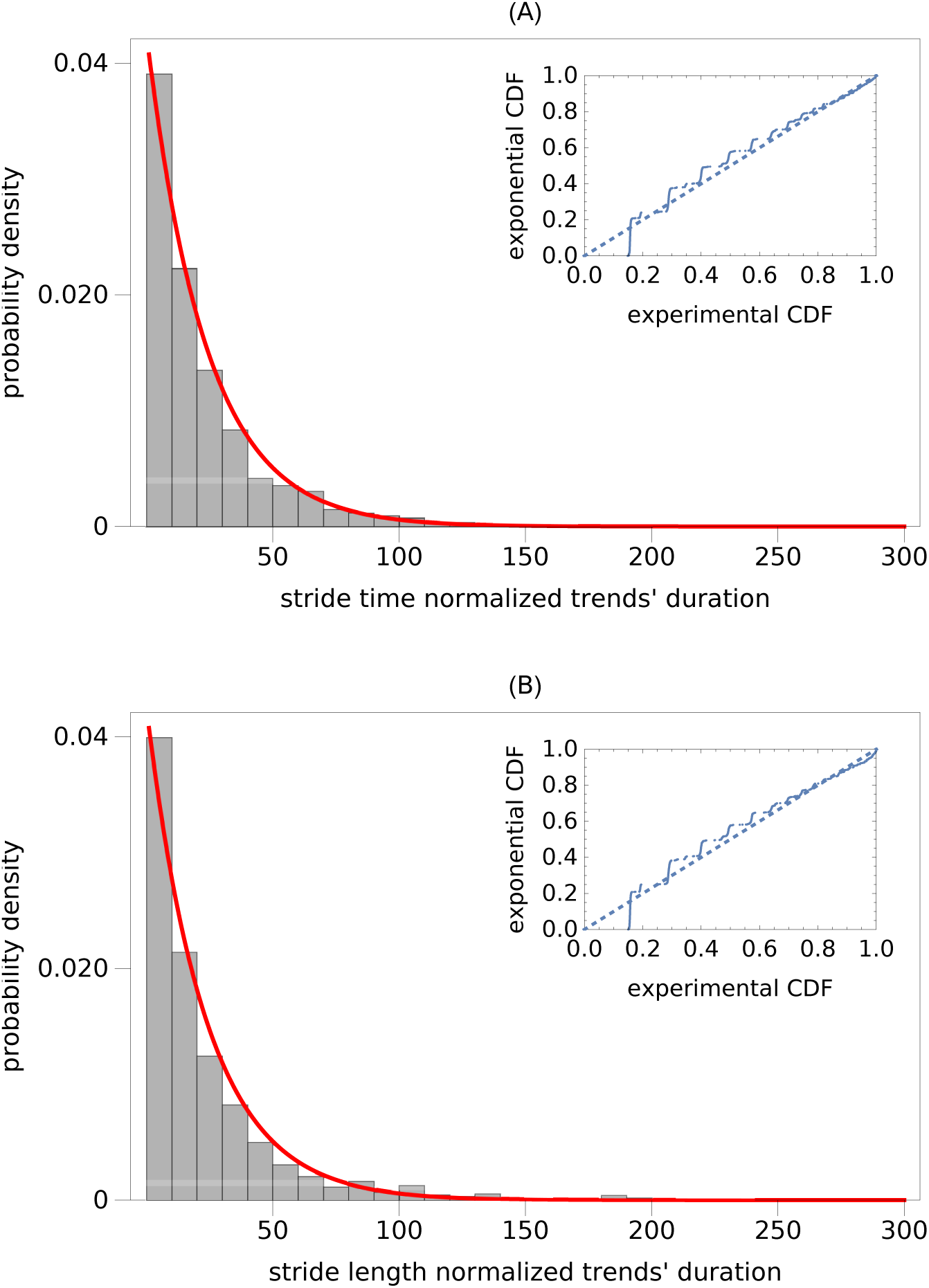
Probability density function (PDF) of normalized duration of trends for: (A) stride time and (B) stride length. Trend durations turned out to be independent of treadmill speed. Consequently, we created two aggregated data sets from trials at different speeds. The solid curves in both subplots show the best-fit exponential PDFs. The probability plots, presented as insets, show the differences between the experimental and exponential cumulative distribution functions (CDFs).

The fundamental property of the exponential distribution, whose probability density function (PDF) is equal to *exp*(−*λx*)*λ*, is that its average value and standard deviation are both equal to 1*/λ*. Therefore, in Fig. 2 we also present the exponential PDFs fitted to the experimental data. For both ST and SL, *λ* = 0.043. It is apparent from the probability plots, presented as insets in Fig. 2, that the exponential distribution fairly approximates the tail of the experimental distributions.

We defined a normalized slope of a trend as the change of gait parameter normalized by the product of average value of this parameter during the trial and the normalized trend duration. The normalized slopes were independent of treadmill’s speed. Consequently, for both ST and SL, we merged the data from all trials. In Fig. 3 we present the PDFs of normalized ST and SL slopes. While the Anderson-Darling test showed that the distributions of ST and SL slopes were different (*p* = 0.04), other statistical tests such as the Kolmogorov-Smirnov (*p* = 0.16) or Cramér-von Mises (*p* = 0.11) indicated just the opposite. The thick lines in this figure are the Cauchy’s PDFs fitted to experimental data. The probability plots indicate fair fits. However, the Anderson-Darling test demonstrated that the ST and SL slopes were not drawn from the Cauchy distribution (in both cases *p* = 6 × 10^−3^). The scale parameter *γ* of the Cauchy distribution was equal to 1.2 × 10^−3^ and 1.5 × 10^−3^ for ST and SL, respectively. The location parameter (location of the maximum of PDF) was equal to 8 × 10^−4^ for ST and 6 × 10^−4^ for SL.

**Fig 3.**
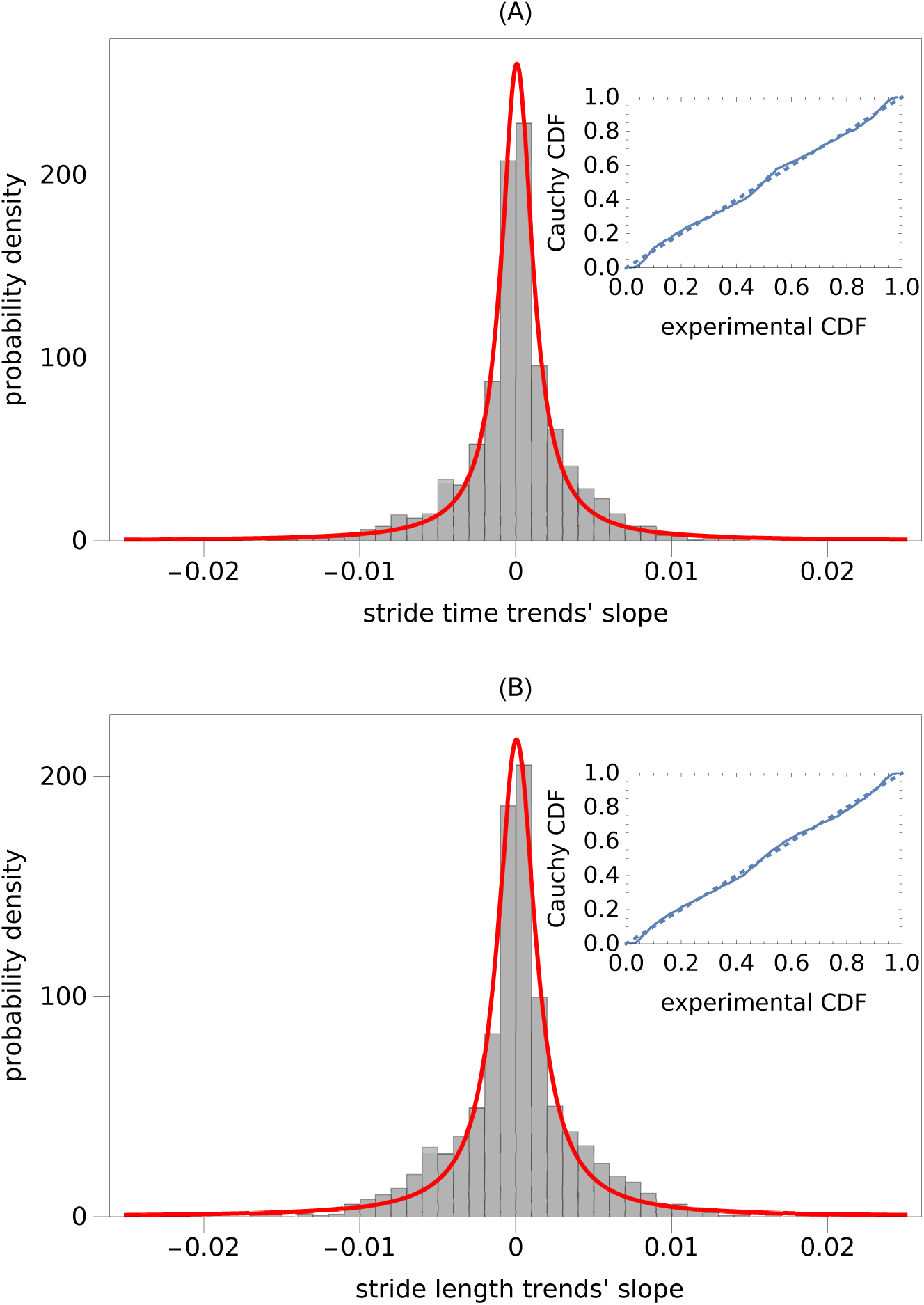
Probability density function (PDF) of normalized slopes of trends in: (A) stride time and (B) stride length. Slopes turned out to be independent of treadmill speed. Consequently, we created two aggregated data sets from trials at different speeds. The solid curves in both subplots show the best-fit Cauchy PDFs. The probability plots, presented as insets, show the differences between the experimental and Cauchy’s cumulative distribution functions (CDFs).

### Scaling exponents

The mean values of ST, SL, and SS scaling exponents at different treadmill speeds are presented in Table 1 along with the corresponding standard deviations. All indices that are statistically smaller than 0.5 are marked with the dagger symbol. The parameters *α*^(1)^, *α*^(2)^, and *α*^(3)^ are exponents calculated using detrended fluctuation analysis of order 1 (DFA1), 2 (DFA2), and 3 (DFA3), respectively. The parameter *α*^(*MD*)^ was computed using the madogram estimator (MD). Scaling analysis was performed for the raw experimental time series and time series detrended using the piecewise linear variant of the MARS model. In the latter case, the exponents have an *L* subscript. Fig. 4 displays the results of the scaling analysis for ST and SL time series.

**Table 1.** Scaling exponents of spatio-temporal gait parameters. *α*^(1)^, *α*^(2)^, and *α*^(3)^ are exponents calculated using detrended fluctuation analysis of order 1 (DFA1), 2 (DFA2), 3 (DFA3), and madogram (MD), respectively. Scaling analysis was performed for the raw experimental time series and time series detrended using the piecewise linear variant of the MARS model. In the latter case the exponents have an *L* subscript. All indices that are statistically smaller than 0.5 are marked with the dagger symbol.

| PWS [%] | $\alpha^{(1)}$ | $\alpha^{(2)}$ | $\alpha^{(3)}$ | $\alpha^{(MD)}$ | $\alpha_L^{(1)}$ | $\alpha_L^{(2)}$ | $\alpha_L^{(3)}$ | $\alpha_L^{(MD)}$ |
| --- | --- | --- | --- | --- | --- | --- | --- | --- |
| **stride length** |  |  |  |  |  |  |  |  |
| 80 | $0.73 \pm 0.12$ | $0.69 \pm 0.12$ | $0.68 \pm 0.13$ | $0.71 \pm 0.10$ | $0.45 \pm 0.07^\dagger$ | $0.49 \pm 0.08$ | $0.52 \pm 0.08$ | $0.49 \pm 0.07$ |
| 90 | $0.70 \pm 0.11$ | $0.67 \pm 0.11$ | $0.66 \pm 0.10$ | $0.67 \pm 0.10$ | $0.49 \pm 0.08$ | $0.52 \pm 0.07$ | $0.54 \pm 0.07$ | $0.49 \pm 0.07$ |
| 100 | $0.77 \pm 0.14$ | $0.70 \pm 0.13$ | $0.68 \pm 0.14$ | $0.69 \pm 0.12$ | $0.46 \pm 0.08^\dagger$ | $0.48 \pm 0.06^\dagger$ | $0.51 \pm 0.07$ | $0.46 \pm 0.06^\dagger$ |
| 110 | $0.75 \pm 0.15$ | $0.70 \pm 0.16$ | $0.67 \pm 0.15$ | $0.69 \pm 0.11$ | $0.43 \pm 0.06^\dagger$ | $0.47 \pm 0.05^\dagger$ | $0.49 \pm 0.05$ | $0.45 \pm 0.05^\dagger$ |
| 120 | $0.76 \pm 0.14$ | $0.71 \pm 0.13$ | $0.69 \pm 0.12$ | $0.71 \pm 0.11$ | $0.44 \pm 0.07^\dagger$ | $0.48 \pm 0.07^\dagger$ | $0.51 \pm 0.07$ | $0.49 \pm 0.07$ |
| all SL | $0.74 \pm 0.13$ | $0.70 \pm 0.13$ | $0.68 \pm 0.13$ | $0.69 \pm 0.11$ | $0.45 \pm 0.07^\dagger$ | $0.49 \pm 0.07^\dagger$ | $0.51 \pm 0.07$ | $0.48 \pm 0.07^\dagger$ |
| **stride time** |  |  |  |  |  |  |  |  |
| 80 | $0.81 \pm 0.16$ | $0.73 \pm 0.16$ | $0.70 \pm 0.16$ | $0.75 \pm 0.12$ | $0.44 \pm 0.07^\dagger$ | $0.47 \pm 0.09^\dagger$ | $0.50 \pm 0.09$ | $0.47 \pm 0.08^\dagger$ |
| 90 | $0.79 \pm 0.11$ | $0.75 \pm 0.13$ | $0.71 \pm 0.13$ | $0.71 \pm 0.12$ | $0.47 \pm 0.08^\dagger$ | $0.51 \pm 0.08$ | $0.54 \pm 0.08$ | $0.48 \pm 0.08$ |
| 100 | $0.85 \pm 0.14$ | $0.78 \pm 0.15$ | $0.76 \pm 0.16$ | $0.75 \pm 0.12$ | $0.45 \pm 0.08^\dagger$ | $0.50 \pm 0.07$ | $0.52 \pm 0.07$ | $0.48 \pm 0.06^\dagger$ |
| 110 | $0.81 \pm 0.13$ | $0.74 \pm 0.16$ | $0.70 \pm 0.16$ | $0.74 \pm 0.12$ | $0.46 \pm 0.07^\dagger$ | $0.50 \pm 0.07$ | $0.52 \pm 0.07$ | $0.49 \pm 0.10$ |
| 120 | $0.82 \pm 0.14$ | $0.74 \pm 0.16$ | $0.72 \pm 0.16$ | $0.75 \pm 0.13$ | $0.45 \pm 0.07^\dagger$ | $0.49 \pm 0.07$ | $0.52 \pm 0.07$ | $0.49 \pm 0.09$ |
| all ST | $0.82 \pm 0.14$ | $0.75 \pm 0.15$ | $0.72 \pm 0.15$ | $0.74 \pm 0.12$ | $0.45 \pm 0.08^\dagger$ | $0.49 \pm 0.08$ | $0.52 \pm 0.08$ | $0.48 \pm 0.08^\dagger$ |
| **stride speed** |  |  |  |  |  |  |  |  |
| 80 | $0.48 \pm 0.07$ | $0.50 \pm 0.08$ | $0.51 \pm 0.09$ | $0.46 \pm 0.07^\dagger$ | $0.47 \pm 0.07^\dagger$ | $0.48 \pm 0.07^\dagger$ | $0.49 \pm 0.08$ | $0.45 \pm 0.07^\dagger$ |
| 90 | $0.47 \pm 0.07^\dagger$ | $0.48 \pm 0.07$ | $0.49 \pm 0.07$ | $0.43 \pm 0.09^\dagger$ | $0.45 \pm 0.06^\dagger$ | $0.47 \pm 0.07^\dagger$ | $0.48 \pm 0.07$ | $0.42 \pm 0.09^\dagger$ |
| 100 | $0.44 \pm 0.08^\dagger$ | $0.44 \pm 0.07^\dagger$ | $0.45 \pm 0.07^\dagger$ | $0.40 \pm 0.09^\dagger$ | $0.42 \pm 0.06^\dagger$ | $0.43 \pm 0.06^\dagger$ | $0.44 \pm 0.06^\dagger$ | $0.39 \pm 0.09^\dagger$ |
| 110 | $0.46 \pm 0.07^\dagger$ | $0.46 \pm 0.08^\dagger$ | $0.47 \pm 0.09$ | $0.40 \pm 0.11^\dagger$ | $0.43 \pm 0.07^\dagger$ | $0.45 \pm 0.08^\dagger$ | $0.46 \pm 0.09^\dagger$ | $0.39 \pm 0.11^\dagger$ |
| 120 | $0.48 \pm 0.07$ | $0.49 \pm 0.09$ | $0.50 \pm 0.09$ | $0.47 \pm 0.10^\dagger$ | $0.45 \pm 0.06^\dagger$ | $0.47 \pm 0.07^\dagger$ | $0.48 \pm 0.08$ | $0.44 \pm 0.08^\dagger$ |
| all SS | $0.47 \pm 0.07^\dagger$ | $0.47 \pm 0.08^\dagger$ | $0.49 \pm 0.09^\dagger$ | $0.43 \pm 0.10^\dagger$ | $0.44 \pm 0.07^\dagger$ | $0.46 \pm 0.07^\dagger$ | $0.47 \pm 0.08^\dagger$ | $0.42 \pm 0.09^\dagger$ |

**Fig 4.**
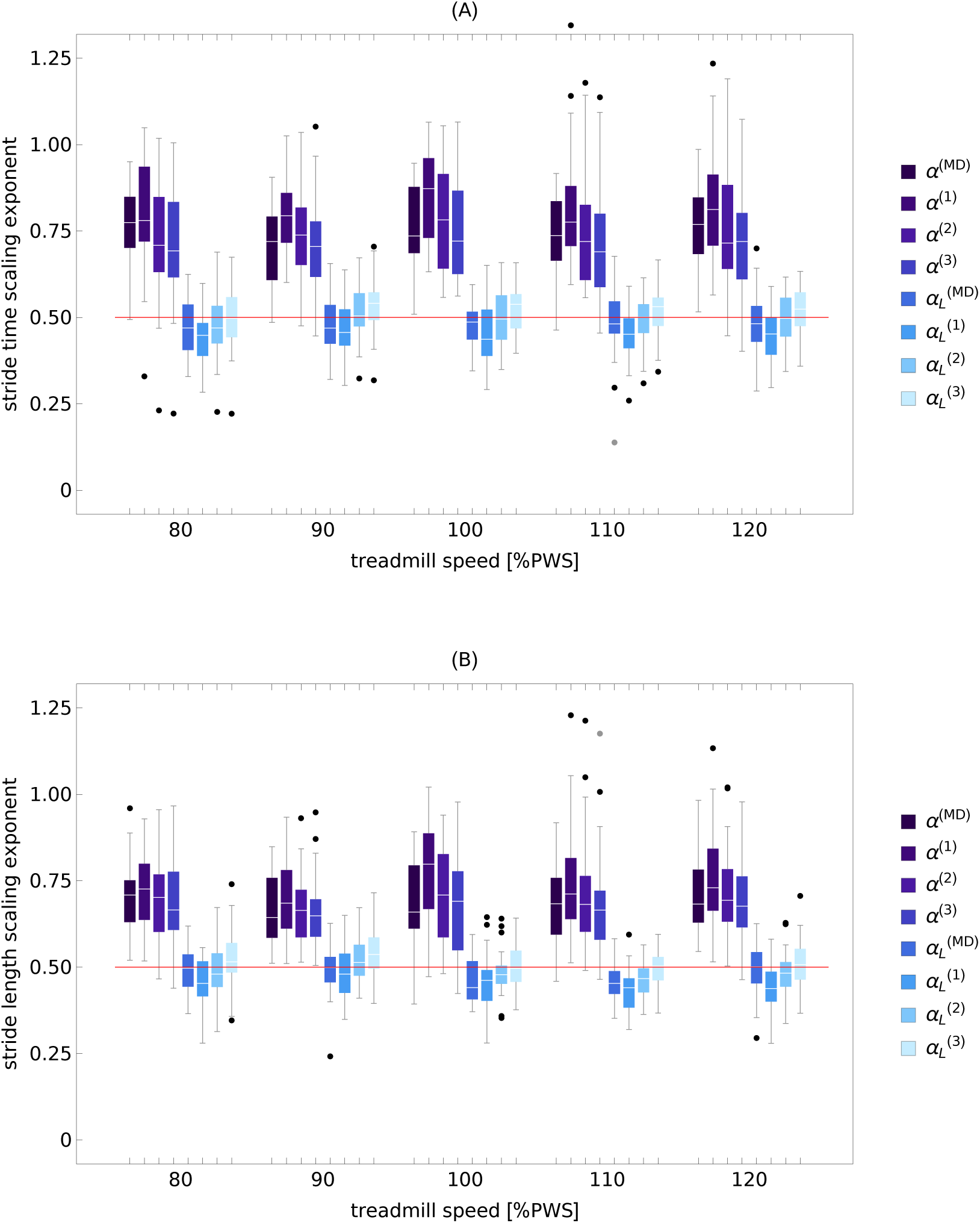
Scaling exponents of stride time (A) and stride length (B) as a function of treadmill speed. *α*^(1)^, *α*^(2)^, *α*^(3)^, and *α*^(*MD*)^ are exponents calculated using detrended fluctuation analysis of order 1 (DFA1), 2 (DFA2), 3 (DFA3), and madogram estimator, respectively. Calculations were performed for raw experimental time series and the time series detrended using the piecewise linear variant of the MARS model. In the latter case the exponents have an *L* subscript. The horizontal red line delineates persistent (*>* 0.5) and anti-persistent (*<* 0.5) scaling.

In the majority of cases, the gait parameters have the following properties:

- The ST, SL, and SS scaling indices do not depend on the treadmill’s speed.
- For all three parameters, there are no statistically significant differences between the scaling exponents *α*^(*n*)^ and *α*^(*MD*)^.
- The scaling exponents of ST and SL MARS residuals are significantly smaller than those of the corresponding experimental time series.

The exceptions are listed below.

For stride speed, the differences in scaling exponents for different speeds were observed for:

- DFA1: 100% PWS vs 120% PWS (*p* = 0.04);
- DFA2: 80% PWS vs 100% PWS (*p* = 0.01), 100% PWS vs 120% PWS (*p* = 0.02);
- DFA3: 80% PWS vs 100% PWS (*p* = 0.02);
- MD: 110% PWS vs 120% PWS (*p* = 0.01), 100% PWS vs 120% PWS (*p* = 0.01).

The scaling exponents *α*^(*n*)^ and *α*^(*MD*)^ were statistically different for:

- 90% PWS: *α*^(1)^ ≠ *α*^(3)^ (*p* = 0.04);
- 90% PWS: *α*^(1)^ ≠ *α*^(*MD*)^ (*p* = 0.02);
- 120% PWS: *α*^(1)^ ≠ *α*^(3)^ (*p* = 0.002).

For stride speed for 90% PWS: *α*^(3)^ ≠ *α*^(*MD*)^ (*p* = 0.02).

To identify the origin of ST and SL persistence, we determined the piecewise linear MARS trends and the corresponding residuals for each trial. We then created an ensemble of 100 composite signals constructed from the original trends and randomly shuffled residuals. We performed DFA*n* and madogram analyses on such ensembles. For both gait parameters and all treadmill speeds, *α*^(*n*)^ and *α*^(*MD*)^ were greater than 0.5. Thus, the outcome of scaling analysis was not determined by the properties of the surrogate noise, which was devoid of any correlations. The results of this numerical experiment for the time series from Fig. 1 are displayed in Fig. 5.

**Fig 5.**
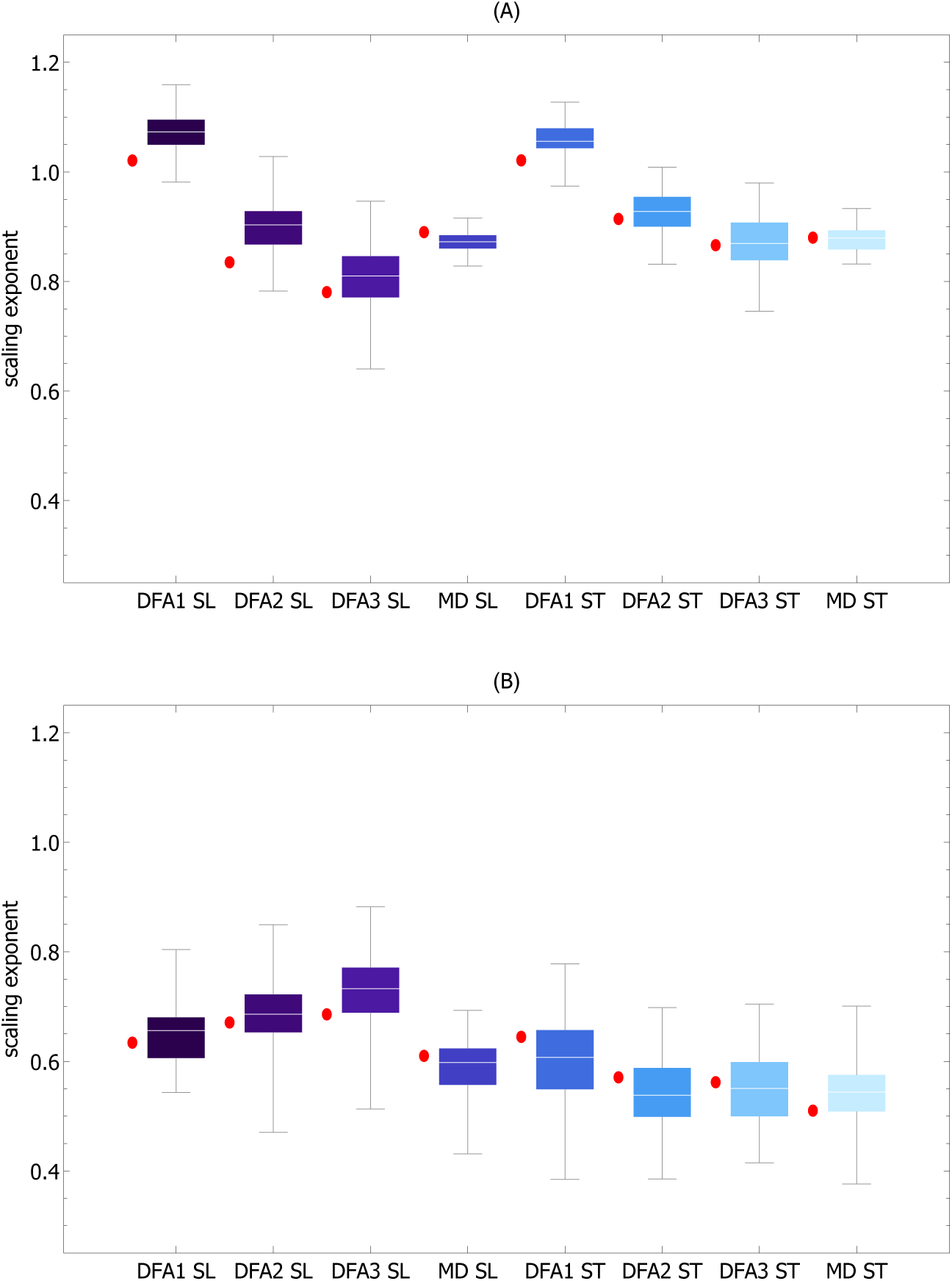
Boxplots of scaling exponents *α*^(*n*)^ and *α*^(*MD*)^ for ensembles of signals comprised of MARS trends extracted from experimental time series and randomly shuffled corresponding MARS residuals. The red disks to the left of each boxplot represent the values of the scaling exponents of the original time series used to generate the ensembles. The calculations were carried out for both stride time (ST) and stride length (SL) time series shown in Fig. 1A and Fig. 1B. Even though there were no correlations in shuffled MARS residuals, the median values of the scaling exponents of composite signals were significantly greater than 0.5 and similar to the scaling exponents of the original time series. Thus, this numerical experiment shows that DFA detrending is incapable of removing trends from ST and SL time series. The values of the scaling exponents of the experimental time series shown in Fig. 1A were: *α*^(1)^ = 1.02, *α*^(2)^ = 0.84, *α*^(3)^ = 0.78, and *α*^(*MD*)^ = 0.89 for SL, and *α*^(1)^ = 1.02, *α*^(2)^ = 0.91, *α*^(3)^ = 0.87, and *α*^(*MD*)^ = 0.88 for ST. For time series in Fig. 1B: *α*^(1)^ = 0.63, *α*^(2)^ = 0.67, *α*^(3)^ = 0.69, and *α*^(*MD*)^ = 0.61 for SL, and *α*^(1)^ = 0.64, *α*^(2)^ = 0.57, *α*^(3)^ = 0.56, and *α*^(*MD*)^ = 0.51 for ST.

### Scaling exponent of short fractional Brownian Motion time series

In Fig. 6 we show the dependence of the scaling (Hurst) exponent of fractional Brownian motion on the length of the data window. Two ensembles of 500 random walks of length 260 with the Hurst exponent equal to *H* = 0.40 and *H* = 0.75 were generated using the Matlab function wfbm. Each trajectory was divided into non-overlapping windows of length *k* from *k* = 40 to *k* = 260 with step 20. For each window, the madogram estimator and detrended fluctuation analysis of order *n* = 1 to *n* = 3 were used to compute *α*^(*MD*)^ and *α*^(*n*)^, respectively. The boxplots of scaling exponents for all four methods are plotted as a function of window length for *H* = 0.40 (Fig. 6A) and *H* = 0.75 (Fig. 6B). These particular values of *H* were chosen because they do not differ significantly from the average value of scaling exponents of MARS residuals and those of ST and SL time series.

**Fig 6.**
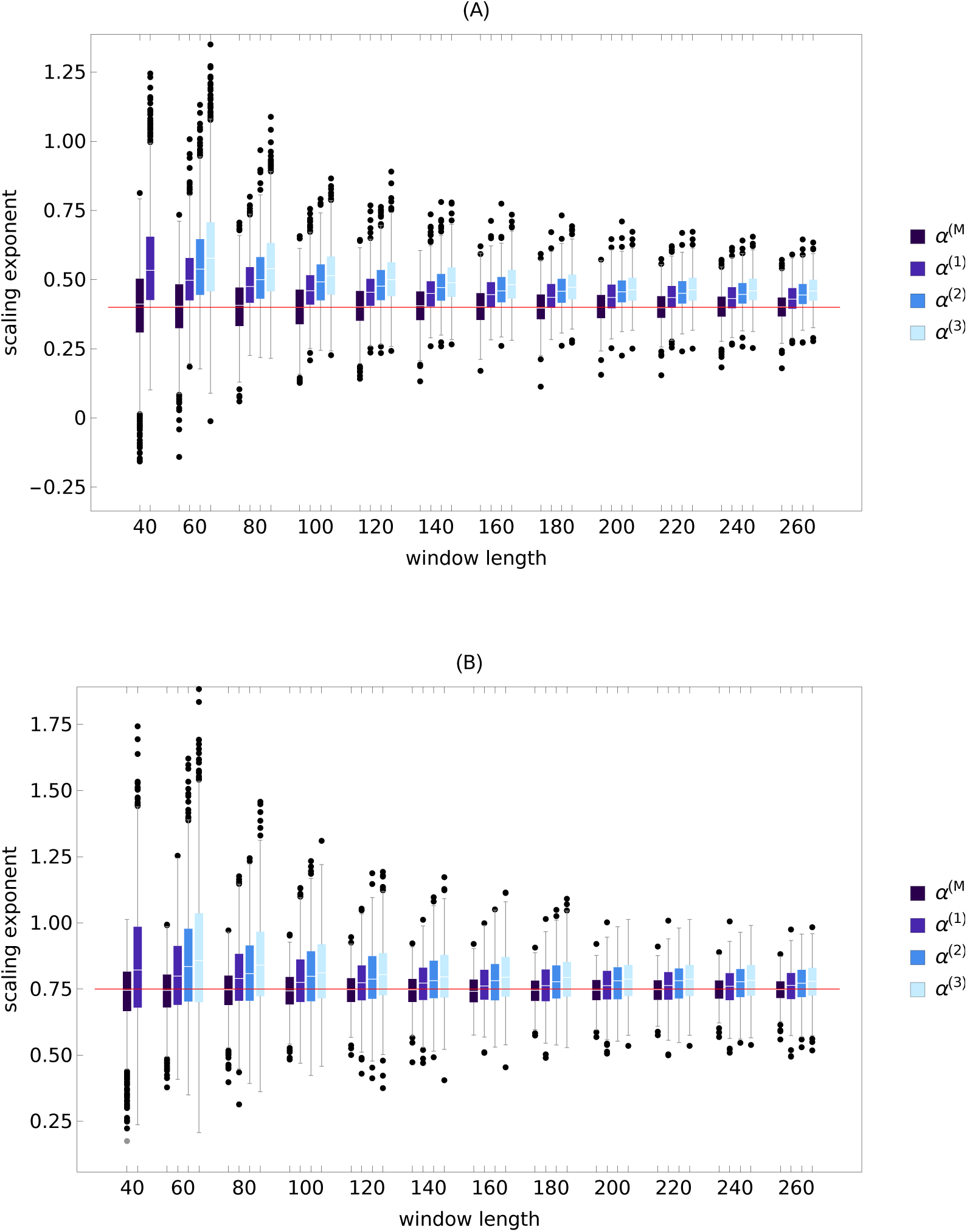
Dependence of scaling exponent of fractional Brownian motion on data window length. Two ensembles of 500 random walks of length 260 with the Hurst exponent equal to *H* = 0.40 an *H* = 0.75 were generated. Each trajectory was divided into non-overlapping windows of length *k* from *k* = 40 to *k* = 260 with step 20. For each window, the madogram estimator and detrended fluctuation analysis of order *n* = 1 to *n* = 3 were used to compute *α*^(*MD*)^ and *α*^(*n*)^, respectively. The boxplots of scaling exponents for all four methods are plotted as a function of window length. Subplot (A) shows the results for *H* = 0.40 and subplot (B) for *H* = 0.75. For the smallest window with *k* = 40, the boxplots for *α*^(2)^ and *α*^(3)^ were not shown due to the large number of outliers.

We can see in Fig. 6 that for short time series DFA*n* overestimates the Hurst exponent. For example, for DFA1 and *k* = 40, the median values were equal to *α*^(1)^ = 0.53 (33% difference) and *α*^(1)^ = 0.82 (9% difference) for *H* = 0.40 an *H* = 0.75, respectively. As window length increases, the estimates of *α*^(*n*)^ converge to the true value. For *k* = 260, the differences dropped to 8% (*α*^(1)^ = 0.43) and 1% (*α*^(1)^ = 0.76), for *H* = 0.40 and *H* = 0.75, respectively. DFA2 and DFA3 were less accurate. For the smallest window, we did not present the boxplots of scaling exponents for these two methods because of the large number of outliers that would obscure Fig. 6. The madogram estimator was the most accurate algorithm. Even for *k* = 40, the estimates were good: *α*^(*MD*)^ = 0.41 and *α*^(*MD*)^ = 0.75 for *H* = 0.40 and *H* = 0.75, respectively. For *H* = 0.40 and the largest window (*k* = 260), the medians of scaling exponents calculated using four methods (madogram and DFA*n*) were different from each other (*p* = 1 × 10^−42^). For *H* = 0.75 and *k* = 260, *α*^(*MD*)^ was significantly smaller than *α*^(*n*)^ for all three orders of detrending. *α*^(1)^ was smaller than *α*^(3)^ (*p* = 7 × 10^−12^).

### Dependence of ST and SL scaling exponents on data window length

Fig. 6 serves as a backdrop for the analysis of analogous calculations performed for stride time and stride length time series. We set the length of the largest window to be smaller than the minimal number of strides observed for the trial at a given treadmill’s speed (such lengths varied from 220 to 260). Consequently, the scaling exponents from all subjects could be included in the analysis. For brevity, we confine the presentation of results to the most accurate algorithm – the madogram estimator.

Fig. 7 shows the dependence of ST and SL scaling exponents on data window length for the 80% PWS experiment. We can see that for *k* = 40 the median of ST scaling scaling exponent *α*^(*MD*)^ = 0.56 is significantly smaller than that for the largest window with *k* = 260 *α*^(*MD*)^ = 0.76 (*p* = 4 × 10^−5^). The same effect is observed for stride length: *α*^(*MD*)^ = 0.58 and *α*^(*MD*)^ = 0.71 for *k* = 40 and *k* = 260, respectively (*p* = 1 × 10^−4^).

**Fig 7.**
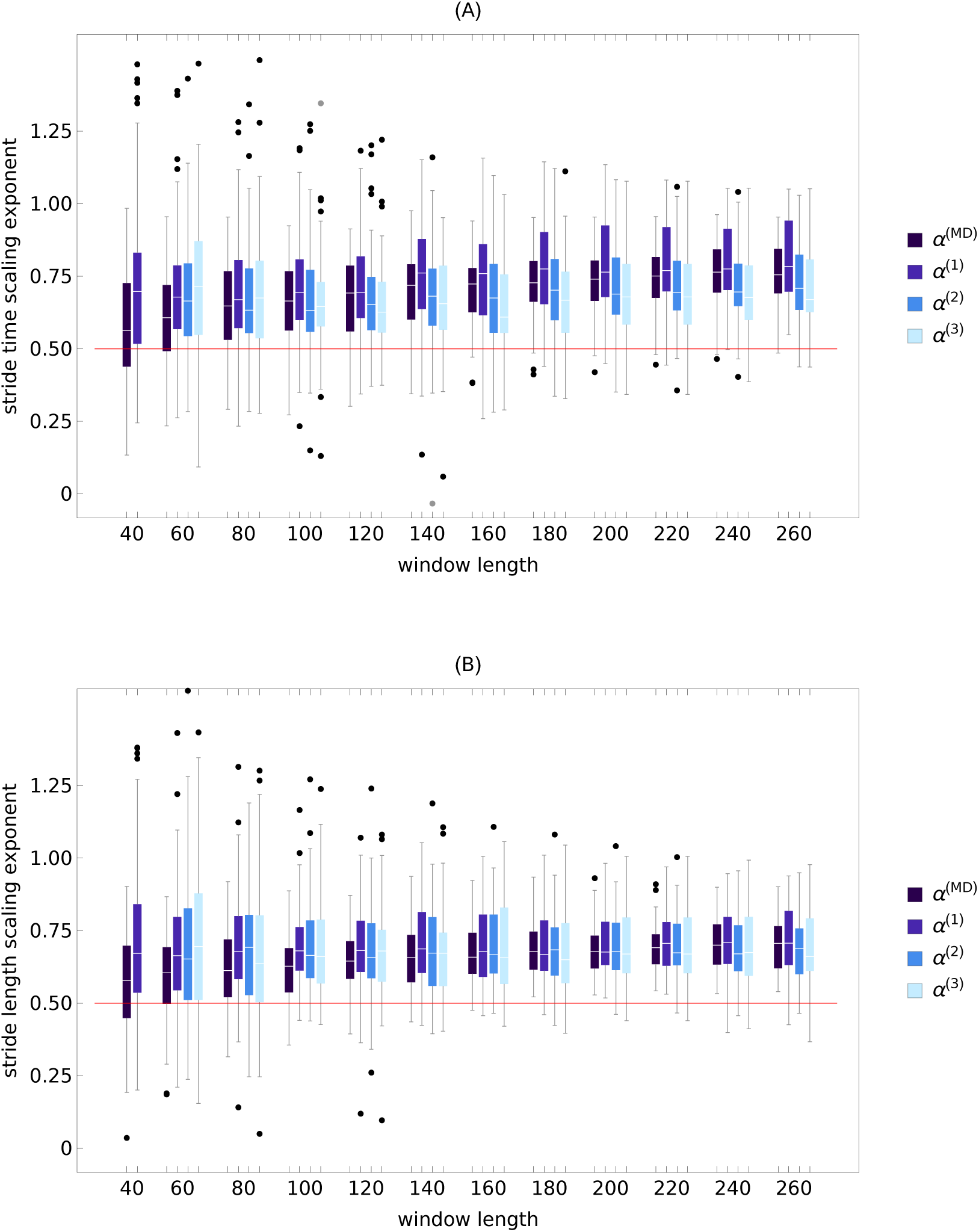
Dependence of scaling exponents of stride time (A) and stride length (B) time series on data window length. Each time series for 80% PWS experiment was divided into non-overlapping windows of length *k* from *k* = 40 to *k* = 260 with step 20. For each window, the madogram estimator and detrended fluctuation analysis of order *n* = 1 to *n* = 3 were used to compute *α*^(*MD*)^ and *α*^(*n*)^, respectively. The boxplots of scaling exponents for all four methods are plotted as a function of window length. For the smallest window with *k* = 40, the boxplots for *α*^(2)^ and *α*^(3)^ were not shown due to the large number of outliers.

In the other four trials, for the smallest window with *k* = 40 the values of *α*^(*MD*)^ for ST were equal to: 0.57, 0.60, 0.62, 0.60 for 90-120% PWS, respectively. For SL, the corresponding values were equal to: 0.57. 0.59, 0.57, 0.59. In all cases, the differences between the smallest and largest window were statistically significant. For *k* = 40, the median of *α*^(*MD*)^ calculated for all five trials was equal to 0.60 and 0.58 for ST and SL, respectively.

### Scaling exponents of parts of gait time series with small slope trends

Fig. 8 presents the boxplots of scaling exponent *α*^(*MD*)^ of the parts of ST and SL time series with small slope trends. More specifically, for each trial we found all segments of MARS trends whose normalized length was greater than 40 and the absolute value of normalized slope was smaller than 0.001 (Fig. 3 shows the distribution of ST and SL slopes). Then, we used the madogram estimator to calculate the scaling exponent of the parts of experimental time series with such trends. The boxplots show the exponents aggregated from all trials (80%-120% PWS). There were 267 values for ST (median 0.57) and 249 values for SL (median 0.56).

**Fig 8.**
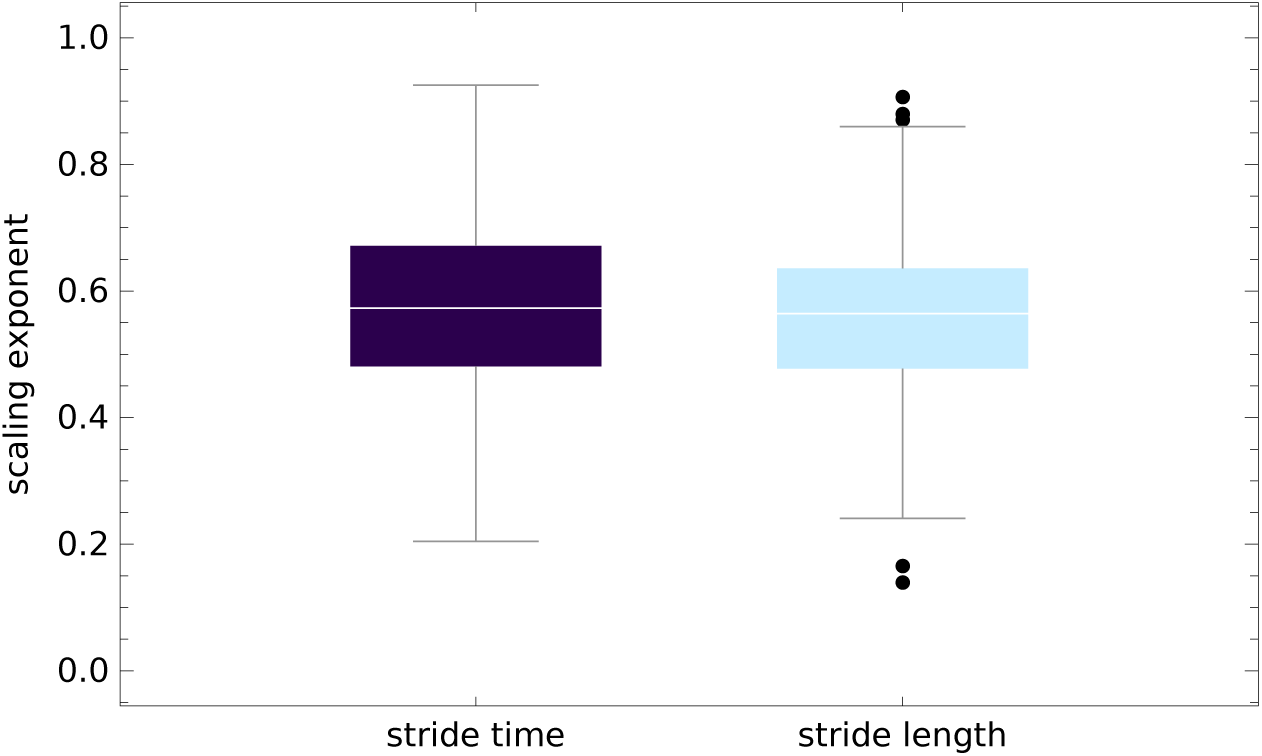
Boxplots of scaling exponent *α*^(*MD*)^ of the parts of stride time (ST) and stride length (SL) series with small slope trends. For each experimental time series, we found all segments of MARS trends whose normalized length was greater than 40 and the absolute value of normalized slope was smaller than 0.001. Then, we extracted the parts of time series with such trends. We used the madogram estimator to calculate the scaling exponent of selected parts. The boxplots show the exponents aggregated from all trials (80%-120% PWS).

In Table 2 we collected the values of *α*^(*MD*)^ for subject 2 for whom in 6 trials the MARS algorithm did not detect linear trends. The median of these values was equal to 0.56.

**Table 2.** The values of *α*^(*MD*)^ for subject 2’s trials with no MARS trends. The scaling exponents are shown for stride length (SL) and stride time (ST).

| parameter | PWS [%] | trial | $\alpha^{(MD)}$ |
| --- | --- | --- | --- |
| SL | 80 | 1 | 0.58 |
| SL | 90 | 1 | 0.61 |
| SL | 100 | 2 | 0.54 |
| SL | 110 | 2 | 0.46 |
| ST | 100 | 2 | 0.60 |
| ST | 110 | 1 | 0.46 |

### Correlation coefficients

Fig. 9 shows the boxplots of Pearson’s correlation coefficient *ρ* between stride time and stride length for all five treadmill speeds. Correlations were calculated for: experimental (raw) time series, trends determined using the MARS algorithm, and MARS residuals (noise). *ρ* was independent of speed: *p*_*raw*_ = 0.12, *p*_*trend*_ = 0.35, and *p*_*noise*_ = 0.17. For the preferred walking speed (PWS), *ρ*_*raw*_ = 0.50 *±* 0.17, *ρ*_*trend*_ = 0.83 *±* 0.12, and *ρ*_*noise*_ = 0.21 *±* 0.10. Thus, coupling between trends was determined to be very strong.

**Fig 9.**
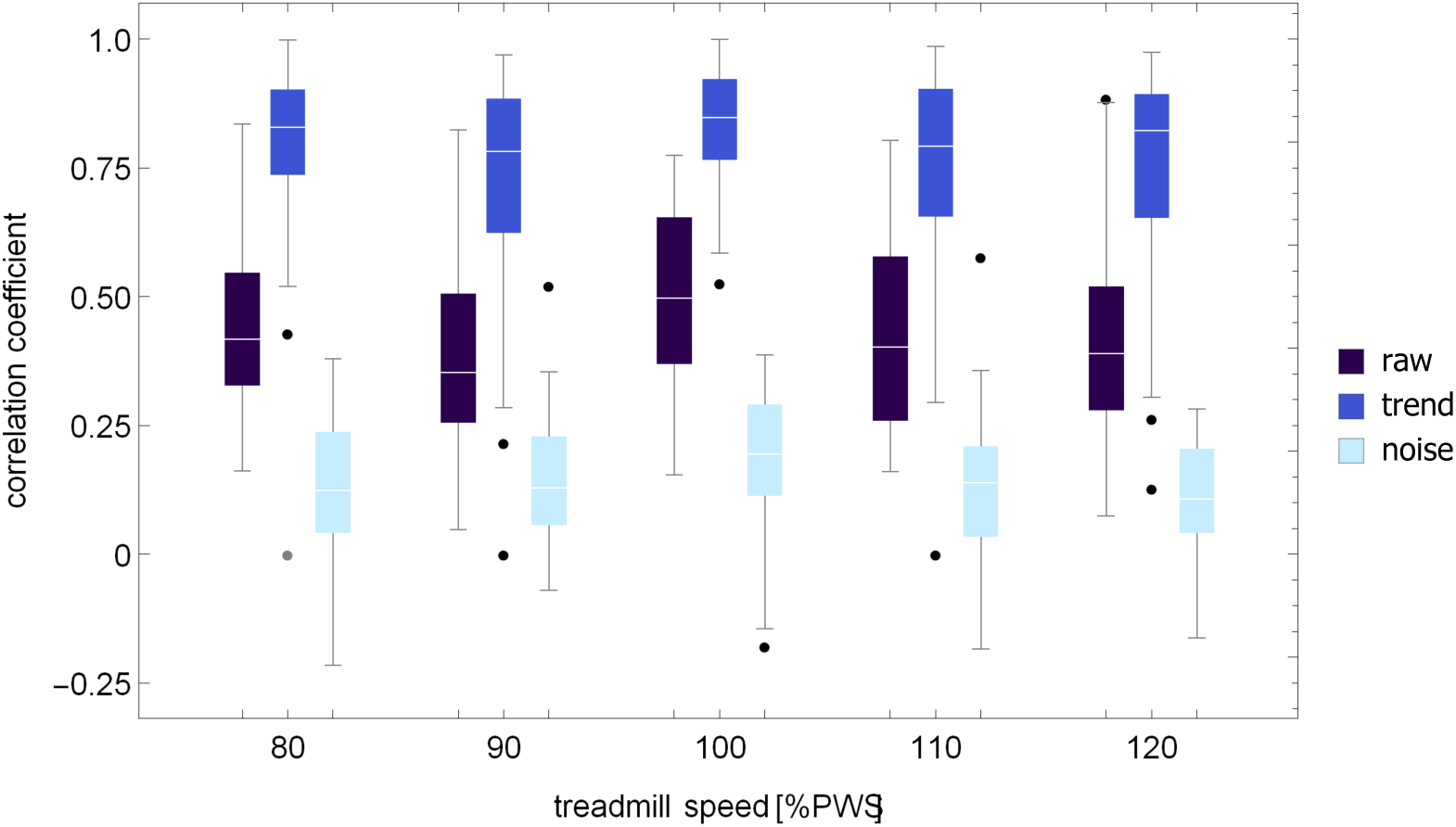
Pearson’s correlation coefficient between stride time and stride length at five different treadmill speeds. Correlations were calculated for: experimental (raw) time series, trends determined using the MARS algorithm, and MARS residuals (noise).

### Trend speed

The examples of time evolution of trend speed *v*^(*trend*)^ are shown in Fig. 1. The boxplots of the trend speed control parameter *TSC* in Fig. 10 show that *v*^(*trend*)^ is tightly controlled about the treadmill speed. For example, at the preferred walking speed, *TSC* = 0.13. For the same speed, the coefficient of variation of *v*^(*trend*)^ was equal to 0.6% (see Fig. 11) and was about three times smaller than that of ST (1.6%), SL (1.8%), and SS (1.7%).

**Fig 10.**
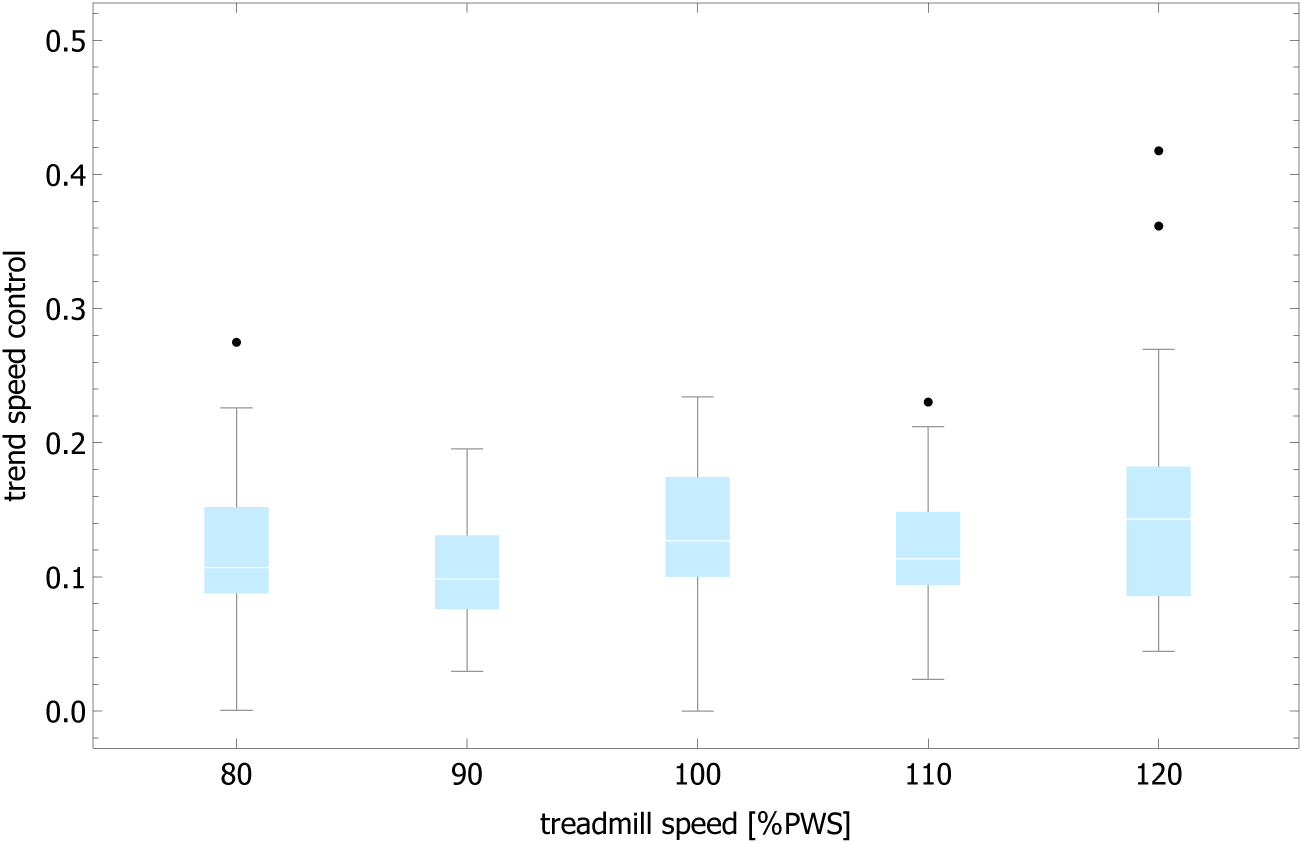
Boxplots of trend speed control parameter (Eq. (6)) for five treadmill speeds.

**Fig 11.**
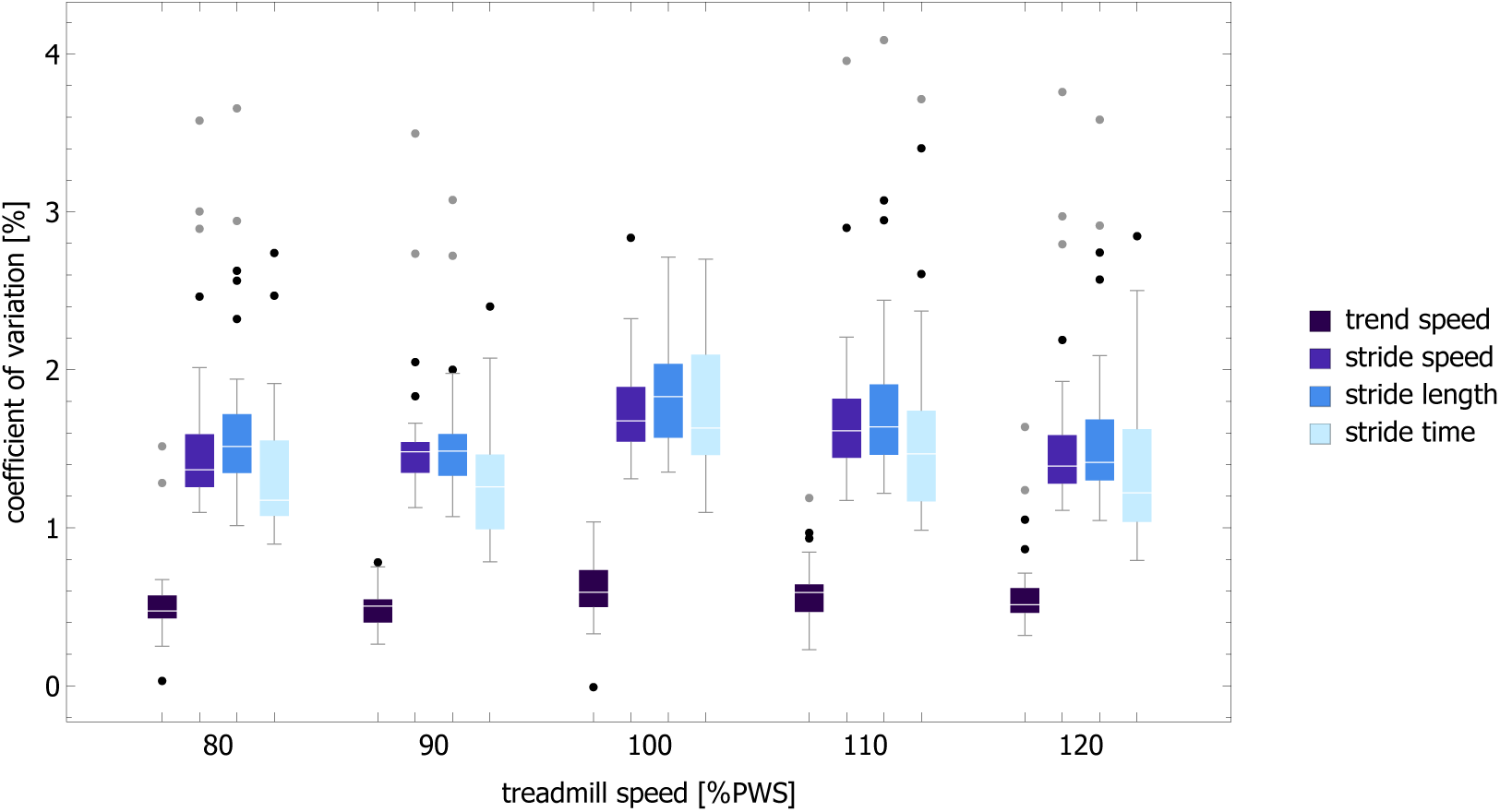
Boxplots of coefficient of variation for trend speed, stride speed, stride length and stride time. The data are shown for five treadmill speeds.

## Discussion

Detrended fluctuation analysis is one of the most frequently used algorithms for fractal analysis of experimental time series. At the time of writing, the seminal paper of Peng et al. [4] has been cited almost 1800 times. It is difficult to overestimate the importance of this method in general and in quantitative gait analysis in particular [1, 2, 14, 16–27].

Early on, numerical investigations showed that simple non-stationarities, such as sinusoidal periodicity or monotonic global trends [28, 29], as well as more complex sources of non-stationarity [30], affected DFA estimates of the scaling exponent. Bryce and Sprague correctly pointed out that, paradoxically, rather than raising doubts about the purported universal applicability of DFA for trended processes, those papers were actually used to demonstrate the efficacy of the method [31].

In DFA, detrending is performed on the integrated signal. Consequently, global linear trends can be eliminated using second or higher order DFA [28, 32]. Figure 1A shows that prominent piecewise linear trends in stride time and stride length may appear even during treadmill walking. Thus, it is surprising that DFA1, which cannot mitigate even global linear trends, was used in most of the previous studies of scaling in spatio-temporal gait parameters. There are two possible explanations for the prevalence of DFA1 in gait analysis. First, while global linear trends distinctly manifest themselves as crossovers in DFA fluctuation function, no apparent crossovers are observed in gait data. Second, the scaling exponents determined using DFA1 and DFA2 are not statistically different, as is evident from the data collected in Table 1, as well as from the visualizations of their distributions in Fig. 4. This agreement was presumably used in favor of DFA1.

We demonstrated that the order of DFA detrending did not have a statistically significant effect on the values of the scaling exponents of the experimental time series. Moreover, *α*^(0)^, *α*^(1)^, and *α*^(2)^ were not significantly different from *α*^(*MD*)^. Thus, we infer that DFA detrending is ineffective, and the observed strong persistence of ST and SL can not stem from the properties of residuals. This interpretation is supported by DFA of signals made up of the MARS trends of experimental time series and randomly shuffled corresponding MARS residuals (shuffled residuals are obviously devoid of any correlations). The fact that *α*^(*n*)^ for both ST and SL were sharply greater than 0.5 (see Fig. 5A) showed that, as in the case of experimental data, strong persistence was associated with the presence of trends.

Let us begin the discussion of significance of trends in gait dynamics with a summary of scaling proprieties of gait parameters. We focus on the results obtained using the most accurate algorithm – the madogram estimator:

- ST and SL scaling exponents *α*^(*MD*)^ increase with the data window length (Fig. 7). For the shortest analyzed window (*k* = 40), the medians of all trials are respectively equal to 0.60 and 0.58. For *k* = 260, fluctuations of gait parameters are strongly persistent (0.75 and 0.71).
- The median of scaling exponent of the parts of the experimental time series with small MARS slopes are equal to 0.57 and and 0.56 for ST and SL, respectively (Fig. 8, see also Table 2).
- Fluctuations of ST and SL MARS residuals are weakly anti-persistent with *α*^(*MD*)^ = 0.48 in both cases (Table 1).

The first two properties strongly indicate that the values of scaling indices of stride time and length are determined by the superposition of large scale trends and small scale fluctuations (note that the fractal dimension of a piecewise linear curve is 1).

We use the second and third property as evidence in support of the hypothesis that trends serve as control manifolds about which ST and SL fluctuate.

Before we discuss the second hypothesis in detail, let us note that during treadmill walking a subject’s stride time and stride length must yield a stride speed that fluctuates over a narrow range centered on the treadmill belt’s speed. In principle, there are infinitely many such combinations which satisfy this type of constraint. These combinations form a straight line in the ST-SL phase space which was dubbed a “Goal Equivalent Manifold” (GEM) [15]. Dingwell et al. projected the deviation vector (calculated in ST-SL space with respect to the mean values of ST and SL) onto the GEM and the axis perpendicular to it. It turned out that fluctuations of tangential and transverse components were persistent and anti-persistent, respectively. Moreover, the tangential variability was higher than the transverse one. Thus, these statistical properties provided the evidence that subjects did not regulate ST and SL independently but instead adjusted them in a coordinated manner to maintain walking speed.

Herein we not only provide *direct* proof of such interdependence but also elucidate how speed control during treadmill walking might actually be accomplished. The distribution of the trend speed control parameter (TSC) in Fig. 10 and coefficient of variation of *v*^(*trend*)^ in Fig. 11 show that the trend speed *v*^(*trend*)^, defined as the ratio of instantaneous values of piecewise linear SL and ST MARS trends, is tightly controlled about the treadmill speed. The strong persistence of stride time and stride length does not impair a subject’s ability to maintain speed because of the strong coupling between their trends (e.g. *ρ*_*trend*_ = 0.85 at PWS). The concomitant changes of instantaneous values of ST and SL trends correspond to movement along the GEM which is the gist of the redundant control of stride speed postulated by Dingwell et al. [33, 34].

We also calculated the correlation coefficient between ST and SL trends as well as the trend speed control parameter using the data from the recent research of Roerdink et al. [35]. While the strength of correlation was comparable to the values presented in this paper, the TSC parameter was smaller, indicating even tighter control of *v*^(*trend*)^. The detailed analysis of Roerdink’s data will be presented elsewhere.

Humans have an innate ability to synchronize their movements with rhythmic sound stimuli. Sejdic et al. argue that sound associated with bipedal gait influenced the evolution of human auditory-motor rhythmic abilities [36]. Walking in sync with a metronome has previously been investigated as a potential rehabilitation tool in patients with Parkinson’s disease [37] and stroke [38]. Isochronous metronome induces anti-persistence in stride time which raises doubts about the efficacy of such rehabilitation. Application of persistent metronomes to gait rehabilitation is an active field of research [20–23, 39].

In this work we demonstrated that the persistence of stride time and length originates from trends that may be approximated by the piecewise linear MARS model. We are not aware of any previous studies that explicitly analyzed the properties of trends in human gait. The normalized trend durations of ST and SL have a distribution with an exponential tail. The exponential distribution describes the time between events in a Poisson point process, i.e. a process in which events occur continuously and independently at a constant average rate. Further research is needed to verify whether this kind of process is involved in the generation of gait trends. We believe that better understanding of gait control mechanisms will pave the way for novel, more effective strategies of gait rehabilitation.

## Conclusions

Hausdorff et al. [1, 2] discovered long-range, persistent correlations in stride duration of human gait nearly a quarter century ago. This persistence was attributed to correlated noise superposed on trends. The evidence presented herein necessitates a fundamental revision of such an interpretation: the statistical properties of ST and SL time series stem from the superposition of large scale trends and small scale fluctuations. We demonstrated that the ST and SL trends serve as control manifolds and that the trend speed is tightly controlled about the treadmill speed. The strong coupling between the ST and SL trends ensures that the concomitant changes of their instantaneous values correspond to movement along the constant speed GEM as postulated by Dingwell et al. [33, 34].

## Materials and methods

### Experimental data

In our analysis, we used data from the study of Dingwell et al. [15], which were available in the Dryad repository [40]. Seventeen young healthy adults were asked to walk on a motor-driven treadmill for five minutes, at five different speeds, determined as a percentage of a subject’s preferred walking speed (PWS). PWS is the speed at which a subject choose to walk on treadmill. There were two trials for each speed (80%, 90%, 100%, 110%, 120% PWS). SL, ST, and SS were determined using a motion capture system. A comprehensive description of the experimental protocol and participants’ characteristics can be found in the original paper.

### Multivariate Adaptive Regression Splines (MARS)

To model trends in spatio-temporal gait parameters we adopted MARS (Multivariate Adaptive Regression Splines) — a nonparametric adaptive regression method proposed by J. Friedman [41, 42]. Let 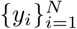 be a time series of experimental values observed at instances 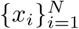. The MARS model *f* is a linear combination of basis functions *h*_*m*_:

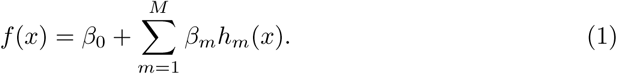

We denote by ℳ the set of basis functions *h*_*m*_ which are constructed in a very specific way. Let us consider two piecewise linear functions *max(0,x*−*t)* and *max(0,t-x)* with a knot at *t*. These two functions (linear splines) form a so-called reflected pair. The example of such a pair is given in Fig 12.

**Fig 12.**
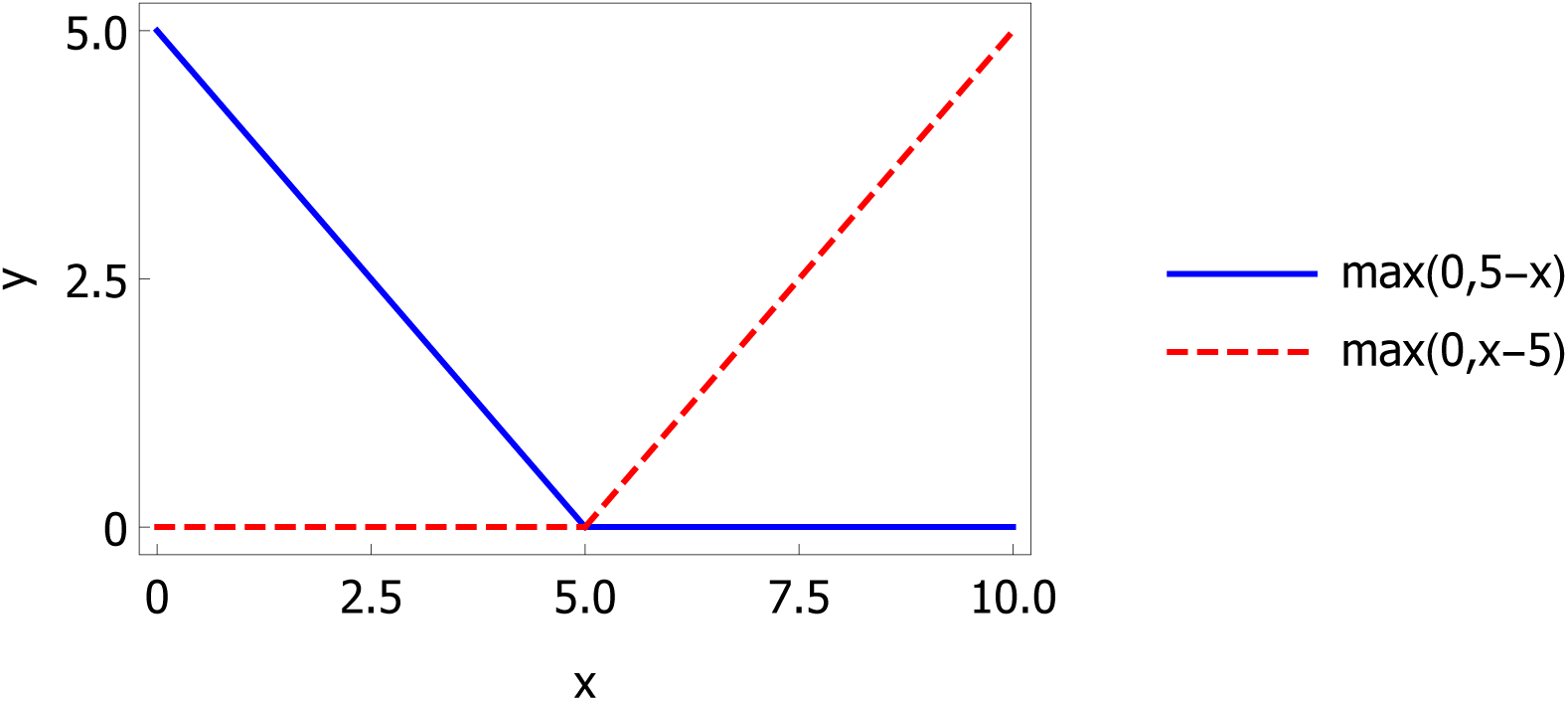
A pair of reflected basis functions with a knot at *t* = 5.

We begin the construction of ℳ by creating a set *C* of *N* reflected pairs with knots equal to values *x*_*i*_:

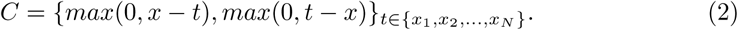

The MARS model is build iteratively. A new basis function *h*_*M*+1_ is the product of one of the already constructed basis functions *h*_*l*_ and one of the reflected pairs from set *C*:

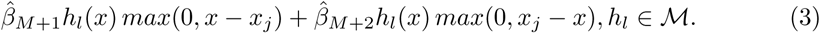

From all possible (*M* + 1)*N* products of this form, we select the one that gives the maximum reduction in sum-of-squares residual error. Here 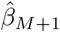 and 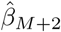 are coefficients estimated by least squares, along with all the other *M* + 1 coefficients in the model. The first basis function *h*_0_(*x*) is constant equal to 1. The process of adding new terms, which is often called a forward model-building procedure (forward pass), is continued until the change in residual error is smaller than the predefined stopping condition or until the maximum number of terms is reached.

The model built during the forward pass is usually overfitted and needs to be pruned to obtain better generalization ability. During each stage of a backward deletion procedure (backward pass), the term whose removal causes the smallest increase in residual squared error is deleted. In this way, we find the best model 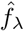 with *λ* terms. The goal of the pruning process is to minimize the value of generalized cross-validation:

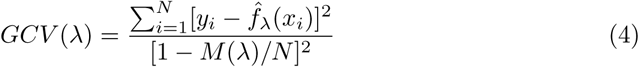

in order to estimate the optimal number of terms. *M* (*λ*) is the effective number of parameters in the model:

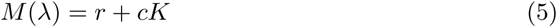

that accounts for both the number of linearly independent basis functions *r* and the number of knots *K* in the forward pass. *c* is a penalty for adding a new knot.

In this study, we used ARESLab, an open source MATLAB implementation of MARS [43], to approximate trends in gait spatiotemporal parameters using a piecewise linear, additive MARS model. In other words, every new basis function was just a linear combination of functions which belonged to a reflected pair (maximum order of interaction was equal to 1). The stopping condition for the forward phase was set to 0.001, and the generalized cross-validation knot penalty *c* was set equal to 2, the value most commonly used for additive models [42].

### Trend speed

We define a trend speed 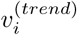 as the ratio of values of piecewise linear SL and ST MARS trends at i*th* stride. To quantify deviations of the trend speed from the average stride speed *< SS >* during a given trial we introduce a trend speed control (TSC) parameter:

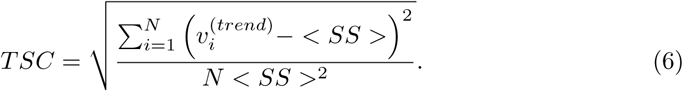

Thus, TSC is equal to zero when the trend and mean stride speed are equal to each other during the whole trial.

### Scaling analysis

#### Detrended fluctuation analysis (DFA)

DFA was designed to detect long-time, power-law scaling of the second moment in the presence of additive, polynomial non-stationarity. Given a bounded time series 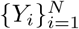, one creates an unbounded process

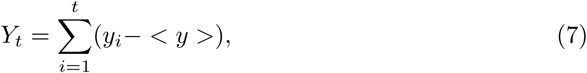

where *< y >* denotes the mean value of the time series. Next, *Y*_*t*_ is divided into windows of length *n* and a local least-squares polynomial fit is performed within each window. Let *P*_*t*_ indicate the piece-wise sequence of such polynomial fits. Then, the root-mean-square deviation from the trend, the fluctuation function, is calculated:

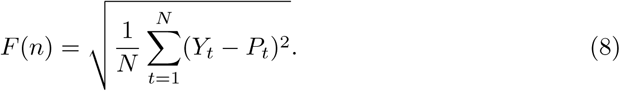

A straight line on a log-log graph of *F* (*n*) as a function of *n* is considered a manifestation of self-affinity as *F* (*n*) ∝ *n*^*α*^. For fractional Gaussian noise, the exponent (index) *α* varies between 0 and 1. In this case, for *α <* 0.5 fluctuations are anti-persistent and for *α >* 0.5 they are persistent. The *n*-th order polynomial regression in the DFA family is typically denoted as DFA*n*. To facilitate reproducibility of the results, we used the Matlab function dfa from the WFDB Toolbox [44, 45] for DFA calculations.

#### Madogram estimator (MD)

The variogram of order *p* of a stochastic process with stationary increments is defined as one half times the expectation value of an increment at lag *t* [46]

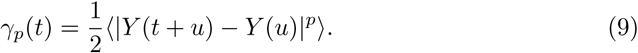

For *p* = 1 we call such a structure function the madogram. As *t* → 0

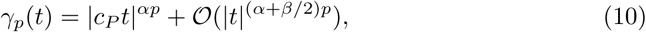

where *H ∈* (0, 1] and constants *β* and *c*_*P*_ are positive. For one dimensional time series 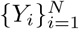, we may define the power variation of order *p*:

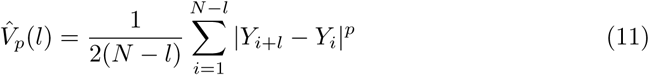

and using Eq. (10) derive the following estimate of fractal dimension:

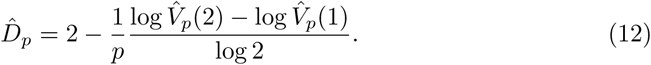

The fractal dimension 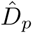 and scaling exponent *α* are related by the following equation:

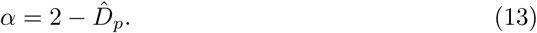

Despite its simplicity, the madogram estimator (*p* = 1) turns out to be particularly robust, especially for non-Gaussian processes. Gneiting er al. [46] compare the properties of fractal dimension estimators that were implemented in the R package fractaldim.

## Statistical analysis

We used the Shapiro-Wilk test to determine whether the analyzed data were normally distributed. The significance threshold for all the statistical tests was set to 0.05.

The dependence of trend durations of stride time and stride length on treadmill speed were examined with the Kruskal-Wallis test with Tukey’s post hoc comparisons.

We compared the values of Pearson’s correlation coefficient between stride time and stride length (experimental values, piecewise linear trends, and MARS residuals) at different speeds using ANOVA or the Kruskal-Wallis test (depending on normality of data). In both cases we used Tukey’s post hoc comparisons.

For a given gait parameter and treadmill speed, the differences in scaling exponents determined using DFA*n* and madogram estimator were assessed with either ANOVA or the Kruskal-Wallis test (with Tukey’s post hoc comparisons in both cases). Such statistical analysis was performed for both original and MARS detrended time series.

To determine whether fluctuations of spatio-temporal gait parameters were anti-persistent, we checked whether the corresponding scaling exponents were smaller than 0.5 using left-sided tests (*t*-test or the Wilcoxon signed rank test).

The Anderson-Darling test was used to compare ST and SL trend duration distributions and to verify whether these distributions were exponential.

In the presented boxplots, the black and grey dots correspond to the outliers and far outliers, respectively.

The analysis of statistical properties of stride time and length trends was carried out using Mathematica 11.3 (Wolfram Research). The other calculations were performed with MATLAB R2016a (Mathworks).

## Author Contributions

Conceptualization: Klaudia Kozlowska, Miroslaw Latka

Formal analysis: Klaudia Kozlowska, Miroslaw Latka, Bruce J. West

Investigation: Klaudia Kozlowska, Miroslaw Latka

Methodology: Klaudia Kozlowska, Miroslaw Latka, Bruce J. West

Software: Klaudia Kozlowska

Supervision: Miroslaw Latka

Validation: Miroslaw Latka

Visualization: Klaudia Kozlowska

Writing – original draft: Miroslaw Latka, Klaudia Kozlowska

Writing – review & editing: Bruce J. West

